## Supplemental figure and tables for "The tuning of tuning: how adaptation influences single cell information transfer"

### Supplementary Figure

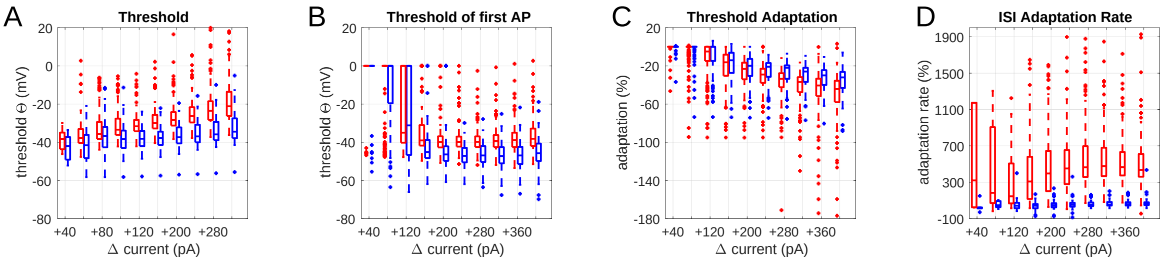


**Fig S1: Threshold behaviour of in the current clamp step-and-hold protocol.** A) Thresholds of all spikes during the step protocol. B) Thresholds of the first spikes after the step current initiation. IC Threshold adaptation: difference in threshold between the first and the last spike of the response. D) Last ISI length relative to the first ISI of the response. Excitatory (red) and inhibitory (blue) neurons. NB Results for significance testing in table S1.

**Supplementary tables**

| Current Injected | Mean AP threshold | First AP threshold | AP threshold Adaptation Rate | Maximum Firing Rate | Latency to first AP |
|---|---|---|---|---|---|
| **+40pA** | d = -0.60<br>CI [-11.2, 3.7]<br>p = 0.30 | d = -0.29<br>CI [-6.3, -0.02]<br>p = 0.048 | d = -0.056<br>CI [-1.7, 1.1]<br>p = 0.70 | d = 0.36<br>CI [0.26, 2.3]<br>p = 0.01 | d = 0.2<br>CI [-101, 136]<br>p = 0.75 |
| **+80pA** | d = -0.60<br>CI [-13.6, -0.15]<br>p = 0.045 | d = -0.15<br>CI [-8.2, 2.5]<br>p = 0.3 | d = -0.075<br>CI [-4.9, 2.7]<br>p = 0.61 | d = 0.71<br>CI [3.8, 9.1]<br>$p = 2.4*10^{-6}$ | d = -0.84<br>CI [-121, -21]<br>p = 0.0062 |
| **+120pA** | d = -0.42<br>CI [-8.5, -0.34]<br>p = 0.034 | d = -0.066<br>CI [-7.3, -4.6]<br>p = 0.65 | d = -0.0055<br>CI [-5.6, 5.3]<br>p = 0.97 | d = 0.90<br>CI [8.1, 15]<br>$p = 3.9*10^{-9}$ | d = -0.75<br>CI [-67, -21]<br>$p = 2.1*10^{-4}$ |
| **+160pA** | d = -0.84<br>CI [-9.2, -4.2]<br>$p = 4.1*10^{-7}$ | d = -0.38<br>CI [-11.1, -1.5]<br>p = 0.0097 | d = 0.070<br>CI [-4.1, 6.8]<br>p = 0.63 | d = 1.4<br>CI [18, 27]<br>$p = 1.0*10^{-19}$ | d = -0.74<br>CI [-44, -18]<br>$p = 6.6*10^{-6}$ |
| **+200pA** | d = -1.1<br>CI [-8.9, -5.2]<br>$p = 2.0*10^{-12}$ | d = -0.65<br>CI [-11.1,-4.3]<br>$p = 1.2*10^{-5}$ | d = 0.16<br>CI [-2.2, 7.7]<br>p = 0.28 | d = 2.2<br>CI [31, 40]<br>$p = 8.0*10^{-38}$ | d = -1.05<br>CI [-28, -15]<br>$p = 3.0*10^{-11}$ |
| **+240pA** | d = -1.2<br>CI [-10.5, -6.4]<br>$p = 9.1*10^{-15}$ | d = -1.2<br>CI [-10.3, -6.2]<br>$p = 8.9*10^{-14}$ | d = 0.32<br>CI [0.5, 9.1]<br>p = 0.028 | d = 3.3<br>CI [42, 51]<br>$p = 4.6*10^{-59}$ | d = -1.03<br>CI [-21, -11]<br>$p = 3.7*10^{-11}$ |
| **+280pA** | d = -1.2<br>CI [-11.4, -7.1]<br>$p = 1.8*10^{-15}$ | d = -1.2<br>CI [-9.3, -5.7]<br>$p = 4.7*10^{-14}$ | d = 0.47<br>CI [3.8, 16.4]<br>p = 0.0017 | d = 4.3<br>CI [51, 58]<br>$p = 4.4*10^{-78}$ | d = -1.4<br>CI [-12.4, -8.3]<br>$p = 3.5*10^{-19}$ |
| **+320pA** | d = -1.3<br>CI [-12.8, -8.1]<br>$p = 1.1*10^{-15}$ | d = -1.2<br>CI [-10.2, -6.2]<br>$p = 1.6*10^{-14}$ | d = 0.30<br>CI [0.46, 30.2]<br>p = 0.043 | d = 4.9<br>CI [58, 65]<br>$p = 2.4*10^{-88}$ | d = -1.5<br>CI [-9.2, -6.2]<br>$p = 5.1*10^{-21}$ |

| | | | | | |
|---|---|---|---|---|---|
| **+360pA** | d = -1.3<br>CI [-14.6, -9.1]<br>p = 2.4*10$^{-15}$ | d = -1.2<br>CI [-11.0, -6.6]<br>p = 1.7*10$^{-13}$ | d = 0.14<br>CI [-9.2, 27.1]<br>p = 0.33 | d = 5.4<br>CI [64, 71]<br>p = 2.02*10$^{-96}$ | d = -1.5<br>CI [-7.1, -4.8]<br>p = 1.3*10$^{-20}$ |
| **+400pA** | d = -1.2<br>CI [-17.1, -10.4]<br>p = 2.7*10$^{-14}$ | d = -1.1<br>CI [-12.0, -7.1]<br>p = 4.9*10$^{-13}$ | d = 0.21<br>CI [-5.0, 33]<br>p = 0.15 | d = 5.6<br>CI [68, 76]<br>p = 1.7*10$^{-99}$ | d = -1.5<br>CI [-5.6, -3.7]<br>p = 5.8*10$^{-20}$ |

| **Current Injected** | **Mean AHP Peak Amplitude** | **Mean AP Half-Width** | **ISI Adaptation Rate** | **ISI Mean** | **ISI Minimum** |
|---|---|---|---|---|---|
| **+40pA** | d = -0.18<br>CI [-11, 3.7]<br>p = 0.29 | d = -1.2<br>CI [-0.70, -0.08]<br>p = 0.016 | d = -0.65<br>CI [-2.1*10$^{3}$, 526]<br>p = 0.7020 | d = -0.75<br>CI [-77, 21]<br>p = 0.23 | d = 0.079<br>CI [-60, 68]<br>p = 0.88 |
| **+80pA** | d = 0.02<br>CI [-13, -0.15]<br>p = 0.045 | d = -1.8<br>CI [-0.86, -0.43]<br>p = 1.7*10$^{-7}$ | d = -0.65<br>CI[-1.1*10$^{3}$, -13]<br>p = 0.044 | d = -1.5<br>CI [-110, -47]<br>p = 8.1*10$^{-6}$ | d = -0.81<br>CI [-84, -11]<br>p = 0.011 |
| **+120pA** | d = 0.04<br>CI [-8.5, -0.34]<br>p = 0.034 | d = -2.02<br>CI [-0.9, -0.6]<br>p = 1.9*10$^{-18}$ | d = -0.63<br>CI [-512, -90]<br>p = 0.0056 | d = -1.2<br>CI [-87, -42]<br>p = 1.3*10$^{-7}$ | d = -0.69<br>CI [-70, -16]<br>p = 0.0016 |
| **+160pA** | d = 0.01<br>CI [-9.2, -4.1]<br>p = 4.1*10$^{-7}$ | d = -2.1<br>CI [-0.94, -0.69]<br>p = 9.2*10$^{-29}$ | d = -1.2<br>CI [-481, -272]<br>p = 3.1*10$^{-11}$ | d = -1.3<br>CI [-64, -38]<br>p = 1.8*10$^{-12}$ | d = -0.53<br>CI [-38, -8.9]<br>p = 0.0018 |
| **+200pA** | d = 0.11<br>CI [-8.9, -5.2]<br>p = 2.0*10$^{-12}$ | d = -2.2<br>CI [-1.04, -0.80]<br>p = 7.0*10$^{-35}$ | d = -1.3<br>CI [-521, -330]<br>p = 6.3*10$^{-16}$ | d = -1.9<br>CI [-61, -44]<br>p = 2.6*10$^{-27}$ | d = -0.57<br>CI [-25, -7.7]<br>p = 2.5*10$^{-4}$ |
| **+240pA** | d = 0.14<br>CI [-10, -6.4]<br>p = 9.1*10$^{-15}$ | d = -2.1<br>CI [-1.1, -0.86]<br>p = 9.1*10$^{-34}$ | d = -1.4<br>CI [-559, -364]<br>p = 1.0*10$^{-17}$ | d = -1.4<br>CI [-53, -35]<br>p = 5.8*10$^{-19}$ | d = -0.29<br>CI [-17, 0.031]<br>p = 0.0508 |

| | | | | | |
|---|---|---|---|---|---|
| **+280pA** | d = 0.24<br>CI [-11, -7.2]<br>p = $1.8*10^{-15}$ | d = -2.0<br>CI [-1.2, -0.95]<br>p = $1.3*10^{-31}$ | d = -1.7<br>CI [-587, -417]<br>p = $8.3*10^{-25}$ | d = -2.5<br>CI [-43, -34]<br>p = $6.6*10^{-43}$ | d = -0.46<br>CI [-6.8, -1.5]<br>p = 0.0021 |
| **+320pA** | d = 0.36<br>CI [-12, 8.1]<br>p = $1.1*10^{-15}$ | d = -1.9<br>CI [-1.4, -1.0]<br>p = $3.0*10^{-29}$ | d = -1.7<br>CI [-591, -419]<br>p = $1.3*10^{-24}$ | d = -2.8<br>CI [-37, -30]<br>p = $2.3*10^{-50}$ | d = -0.59<br>CI [-4.5, -1.6]<br>p = $7.3*10^{-5}$ |
| **+360pA** | d = 0.49<br>CI [-14.6, -9.1]<br>p = $2.3*10^{-15}$ | d = -1.9<br>CI [-1.5, -1.1]<br>p = $1.4*10^{-28}$ | d = -1.7<br>CI [-570, -406]<br>p = $2.9*10^{-25}$ | d = -3.1<br>CI [-34, -29]<br>p = $2.7*10^{-56}$ | d = -0.77<br>CI [-3.9, -1.8]<br>p = $3.9*10^{-7}$ |
| **+400pA** | d = 0.62<br>CI [-17, -10]<br>p = $2.7*10^{-14}$ | d = -1.8<br>CI [-1.7, -1.3]<br>p = $2.7*10^{-26}$ | d = -1.7<br>CI [-534, -376]<br>p = $7.9*10^{-24}$ | d = -3.1<br>CI [-32, -26]<br>p = $9.9*10^{-55}$ | d = -0.91<br>CI [-3.9, -2.0]<br>p = $3.7*10^{-9}$ |

P = p-value (according to 2-sample t-test)

d = Cohen’s d (Measure of effect power); positive means inhibitory has a higher value.

CI = 95% Confidence Interval (for effect power)

**Supplementary Table S1**: Statistical tests of the comparison between excitatory and inhibitory neurons in the current clamp step protocol.

| | N inh | N exc | mean inh | mean exc | h KS test | p KS test | KS stat | h WR test | p WR test | Cliff's Delta |
|---|---|---|---|---|---|---|---|---|---|---|
| # spikes per up state | 1404 | 560 | 0.99 | 1.22 | 0 | 0.047 | 0.068 | 0 | 0.59 | 2.52 |
| # spikes per down state | 1404 | 560 | 0.15 | 0.49 | 1 | 5.8e-10 | 0.16 | 1 | 9.4e-10 | 2.33 |
| firing rate up (Hz) | 1404 | 560 | 12.8 | 3.52 | 1 | 7e-155 | 0.66 | 1 | 7e-163 | 3.29 |
| firing rate down (Hz) | 1404 | 560 | 1.04 | 0.63 | 1 | 1.4e-43 | 0.35 | 1 | 8.1e-48 | 2.93 |
| normalized firing rate up | 1404 | 560 | 0.64 | 0.79 | 1 | 0.0015 | 0.094 | 0 | 0.0579 | 2.45 |
| normalized firing rate down | 1404 | 560 | 0.052 | 0.138 | 1 | 3.0e-8 | 0.149 | 1 | 1.9e-8 | 2.34 |

**Supplementary Table S2**: Statistical tests of the comparison between excitatory and inhibitory neurons in the frozen noise protocol (see main text Fig. 5). P-values were compared to a threshold of 5% / 6 groups = 0.83 % (Bonferroni correction).

| | N inh control | mean inh control | h KS test | p KS test | KS stat | h WR test | p WR test | Cliff's Delta |
|---|---|---|---|---|---|---|---|---|
| # spikes per up state | 33 | 3.77 | 1 | 1.0e-18 | 0.80 | 1 | 2.5e-16 | 0.91 |
| # spikes per down state | 33 | 2.62 | 1 | 1.0e-19 | 0.82 | 1 | 3.0e-16 | 0.83 |
| firing rate up (Hz) | 33 | 10.87 | 1 | 1.5e-18 | 0.79 | 1 | 2.5e-16 | 0.91 |
| firing rate down (Hz) | 33 | 3.43 | 1 | 1.0e-19 | 0.82 | 1 | 7.3e-16 | 0.89 |
| normalized firing rate up | 33 | 2.62 | 1 | 1.5e-18 | 0.80 | 1 | 2.5e-16 | 0.91 |
| normalized firing rate down | 33 | 0.86 | 1 | 1.0e-19 | 0.82 | 1 | 7.3e-16 | 0.89 |

**Supplementary Table S3**: Statistical tests of the comparison between excitatory and inhibitory neurons receiving the control frozen noise stimulus (see main text Fig. 5). P-values were compared to a threshold of 5% / 6 groups = 0.83 % (Bonferroni correction).
